# Multi-omic rejuvenation of human cells by maturation phase transient reprogramming

**DOI:** 10.1101/2021.01.15.426786

**Authors:** Diljeet Gill, Aled Parry, Fátima Santos, Irene Hernando-Herraez, Thomas M. Stubbs, Inês Milagre, Wolf Reik

**Author notes:** Correspondence (I.M.), (W.R.).

## Abstract

Ageing is the gradual decline in organismal fitness that occurs over time leading to tissue dysfunction and disease. At the cellular level, ageing is associated with reduced function, altered gene expression and a perturbed epigenome. Somatic cell reprogramming, the process of converting somatic cells to induced pluripotent stem cells (iPSCs), can reverse these age-associated changes. However, during iPSC reprogramming somatic cell identity is lost, and can be difficult to reacquire as re-differentiated iPSCs often resemble foetal rather than mature adult cells. Recent work has demonstrated that the epigenome is already rejuvenated by the maturation phase of reprogramming, which suggests full iPSC reprogramming is not required to reverse ageing of somatic cells. Here we have developed the first “maturation phase transient reprogramming” (MPTR) method, where reprogramming factors are expressed until this rejuvenation point followed by withdrawal of their induction. Using dermal fibroblasts from middle age donors, we found that cells reacquire their fibroblast identity following MPTR, possibly as a result of persisting epigenetic memory at enhancers. Excitingly, our method substantially rejuvenated multiple cellular attributes including the transcriptome, which was rejuvenated by around 30 years as measured by a novel transcriptome clock. The epigenome, including H3K9me3 histone methylation levels and the DNA methylation ageing clock, was rejuvenated to a similar extent. The magnitude of rejuvenation instigated by MTPR is substantially greater than that achieved in previous transient reprogramming protocols. MPTR fibroblasts produced youthful levels of collagen proteins, suggesting functional rejuvenation. Overall, our work demonstrates that it is possible to separate rejuvenation from pluripotency reprogramming, which should facilitate the discovery of novel anti-ageing genes and therapies.

**Highlights:**

- We developed a novel method by which human fibroblasts are reprogrammed until the maturation phase of iPSCs and are then returned to fibroblast identity
- DNA methylation memory in fibroblast enhancers may allow recovery of cell identity when fibroblast gene expression programmes are already extinct
- Molecular measures of ageing including transcriptome and DNA methylation clocks and H3K9me3 levels reveal robust and substantial rejuvenation
- Functional rejuvenation of fibroblasts by MPTR is suggested by reacquisition of youthful levels of collagen proteins

## Introduction

Ageing is the gradual decline in cell and tissue function over time that occurs in almost all organisms, and is associated with a variety of molecular hallmarks such as telomere attrition, genetic instability, epigenetic and transcriptional alterations and an accumulation of misfolded proteins (López-Otín et al., 2013). This leads to perturbed nutrient sensing, mitochondrial dysfunction and increased incidence of cellular senescence, which impact overall cell function, intercellular communication, promotes exhaustion of stem cell pools and causes tissue dysfunction (López-Otín et al., 2013). The progression of some ageing related changes, such as transcriptomic and epigenetic ones, can be measured highly accurately and as such they can be used to construct “ageing clocks” that predict chronological age with high precision in humans (Fleischer et al., 2018; Hannum et al., 2013; Horvath, 2013; Peters et al., 2015) and in other mammals (Stubbs et al., 2017; Thompson et al., 2017, 2018). Since transcriptomic and epigenetic changes are reversible at least in principle, this raises the intriguing question of whether molecular attributes of ageing can be reversed and cells phenotypically rejuvenated (Rando and Chang, 2012).

Induced pluripotent stem cell (iPSC) reprogramming is the process by which almost any somatic cell can be converted into an embryonic stem cell like state. Intriguingly, iPSC reprogramming reverses many age-associated changes including telomere attrition and oxidative stress (Lapasset et al., 2011). Notably, the epigenetic clock is reset back to approximately 0, suggesting reprogramming can reverse ageing associated epigenetic alterations (Horvath, 2013). However, iPSC reprogramming also results in the loss of original cell identity and therefore function. By contrast transient reprogramming approaches where Yamanaka factors (Oct4, Sox2, Klf4, c-Myc) are expressed for short periods of time may be able to achieve rejuvenation without loss of cell identity. Reprogramming can be performed *in vivo* (Abad et al., 2013) and indeed, cyclical expression of the Yamanaka factors *in vivo* can extend lifespan in progeroid mice and improves cellular function in wild type mice (Ocampo et al., 2016). An alternative approach for reprogramming *in vivo* demonstrated reversal of ageing-associated changes in retinal ganglion cells and was capable of restoring vision in a glaucoma mouse model (Lu et al., 2020). More recently, *in vitro* transient reprogramming has been shown to reverse multiple aspects of ageing in human fibroblasts and chondrocytes (Sarkar et al., 2020). Nevertheless, the extent of epigenetic rejuvenation achieved by previous transient reprogramming methods has been modest (∼3 years) compared to the drastic reduction achieved by complete iPSC reprogramming. Here we establish a novel transient reprogramming strategy where Yamanaka factors are expressed until the maturation phase of reprogramming before abolishing their induction (maturation phase transient reprogramming, MPTR), with which we were able to achieve robust and very substantial rejuvenation (∼30 years) whilst retaining original cell identity.

## Results

### Transiently reprogrammed cells reacquire their initial cell identity

Reprogramming can be divided into three phases: initiation, maturation and stabilisation (Samavarchi-Tehrani et al., 2010) (Figure 1A). Previous attempts at transient reprogramming have only reprogrammed within the initiation phase (Ocampo et al., 2016; Sarkar et al., 2020). However, reprogramming further, up to the maturation phase, may achieve more substantial rejuvenation. To investigate the potential of maturation-phase transient reprogramming (MPTR) to reverse ageing phenotypes, we generated a doxycycline inducible polycistronic reprogramming cassette that encoded *Oct4, Sox2, Klf4, c-Myc* and GFP. By using a polycistronic cassette, we could ensure that individual cells were able to express all four Yamanaka factors. This reprogramming cassette was capable of generating doxycycline-independent iPSC lines from human fibroblasts and induced a substantial reduction of DNA methylation age throughout the reprogramming process, consistent with previous work using a different reprogramming system (Olova et al., 2018)(Figure 1A). Specifically, DNA methylation age as measured using the epigenetic clock (Horvath, 2013) was substantially reduced relatively early in the reprogramming process (which takes about 50 days to be complete), with an approximate rejuvenation of 20 years by day 10 and 40 years by day 17 (Figure 1A). Similar results were obtained using the skin and blood clock (Horvath et al., 2018) (Supplementary Figure 1A). We therefore focussed on the window between days 10 and 17 to develop our MPTR protocol for human fibroblasts (Figure 1B), predicting that this would enable substantial reversal of ageing phenotypes whilst potentially allowing cells to regain original cell identity. The reprogramming cassette was introduced into fibroblasts from three middle aged donors (38, 53 and 53 years old) by lentiviral transduction before selecting successfully transduced cells by sorting for GFP. We then reprogrammed the fibroblasts for different lengths of time (10, 13, 15 or 17 days) by supplementing the media with 2 µg/mL doxycycline and carried out flow sorting to isolate cells that were successfully reprogramming (labelled “transient reprogramming intermediate”: SSEA4 positive, CD13 negative) as well as the cells that had failed to reprogram (labelled “failing to transiently reprogram intermediate”: CD13 positive, SSEA4 negative). At this stage, approximately 25% of the cells were successfully reprogramming and approximately 35% of the cells were failing to reprogram, whilst the remainder were double positive or double negative (Supplementary Figure 1B). Cells were harvested for DNA methylation array or RNA-seq and also re-plated for further culture in the absence of doxycycline to stop the expression of the reprogramming cassette. Further culture in the absence of doxycycline generated “transiently reprogrammed fibroblasts”, which had previously expressed SSEA4 at the intermediate stage, as well as “failed to transiently reprogram fibroblasts”, which had expressed the reprogramming cassette (were GFP positive) but failed to express SSEA4. As a negative control, we simultaneously ‘mock infected’ populations of fibroblasts from the same donors. These cells underwent an initial flow sort for viability (to account for the effects of the GFP sort) before culture under the same conditions as the reprogramming cells and flow sorting for CD13 (cells harvested at this stage generated a “negative control intermediate” for methylome and transcriptome analyses). Finally, these “negative control intermediate” cells were grown in the absence of doxycycline for the same length of time as experimental samples to account for the effects of extended cell culture, generating “negative control fibroblasts” (Figure 1B).

**Figure 1.**
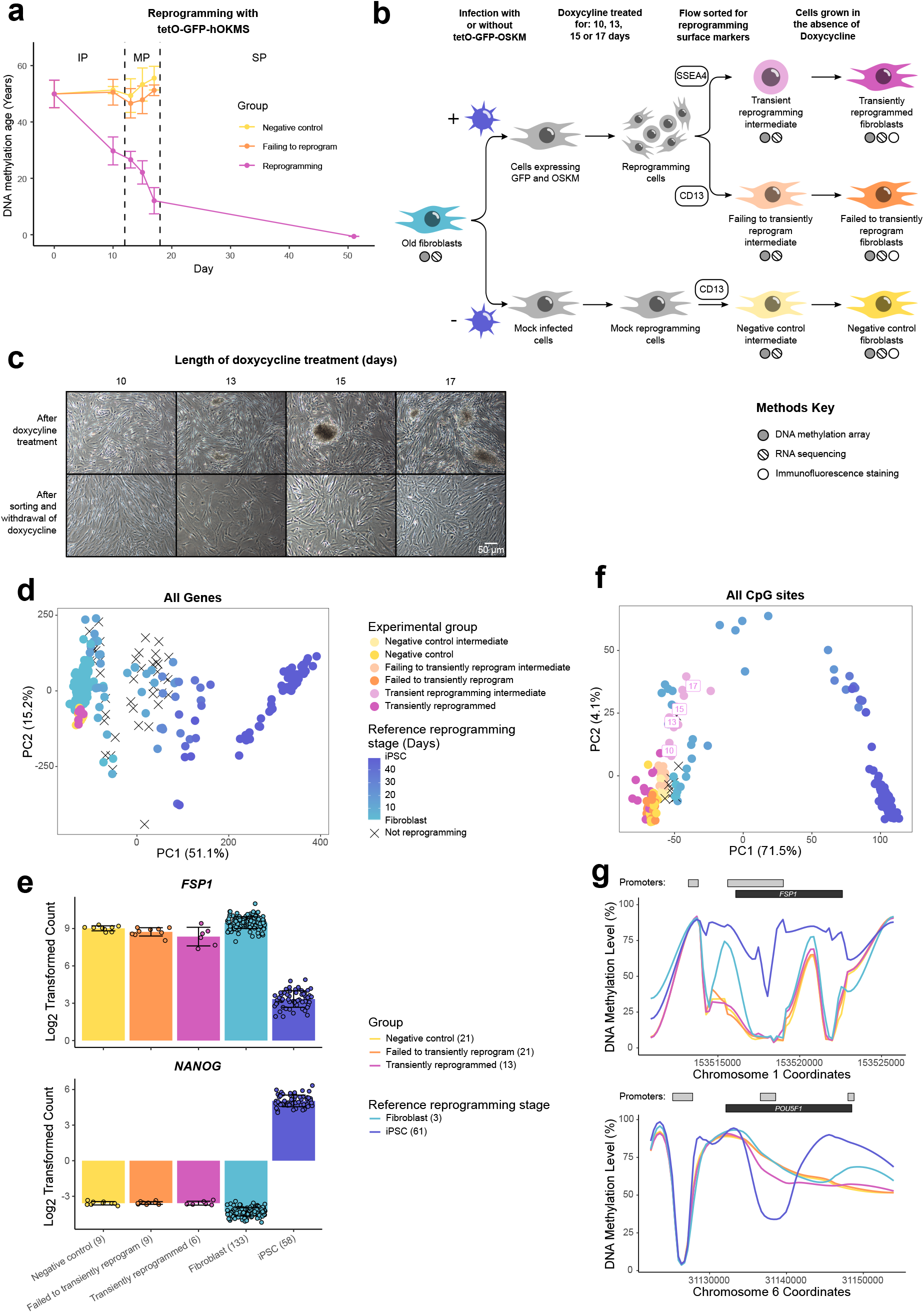
Transiently reprogrammed cells reacquire their initial cellular identity. (A) Mean DNA methylation age (calculated using the multi-tissue clock (Horvath, 2013)) throughout the reprogramming process where cells were transduced with our tetO-GFP-hOKMS vector and treated continuously with 2 µg/mL of doxycycline. Reprogramming is divided in three distinct phases: initiation phase (IP); maturation phase (MP) and stabilisation phase (SP). DNA methylation age decreased substantially during the maturation phase of reprogramming in cells that were successfully reprogramming (magenta line) but not in control cells (yellow and orange lines represent non-transduced cells and cells expressing hOKMS but failing to reprogram as indicated by cell surface markers, respectively). Points represent the mean and error bars the standard deviation. N = 3 biological replicates per condition, where fibroblasts were derived from different donors. N = 2 biological replicates for the iPSC timepoint (day 51). (B) Experimental scheme for maturation phase transient reprogramming (MPTR). The tetO-GFP-hOKMS reprogramming construct was introduced into fibroblasts from older donors by lentiviral transduction. Alternatively, cells were ‘mock infected’ as a negative control. Following this, cells were grown in the presence of 2 µg/mL doxycycline to initiate reprogramming. At several timepoints during the maturation phase, cells were flow sorted and successfully reprogramming cells (CD13-SSEA4+) and cells that were failing to reprogram (CD13+ SSEA4-) were collected for analysis. These were termed “transient reprogramming intermediate” and “failing to transiently reprogram intermediate”, respectively. Sorted cells were also further cultured, and grown in the absence of doxycycline for at least four weeks - these were termed “transiently reprogrammed” (CD13-SSEA4+) or “failed to transiently reprogram” (CD13+ SSEA4-). (C) Phase-contrast microscope images of cells after doxycycline treatment (transient reprogramming intermediate) and after withdrawal of doxycycline (transiently reprogrammed) as described in B. The morphology of some cells changed after doxycycline treatment. These cells appeared to form colonies, which became larger with longer exposure to doxycycline. After sorting, these cells were cultured in medium no longer containing doxycycline, and appeared to return to their initial fibroblast morphology. (D) Principal component analysis of transient reprogramming and reference reprogramming sample transcriptomes (light blue to dark blue, data from Banovich et al (2018), Fleischer et al (2018) and our novel Sendai reprogramming dataset). Reference samples form a reprogramming trajectory along PC1. Transiently reprogrammed samples and control samples were located at the beginning of this trajectory. (E) Mean gene expression levels for the fibroblast specific gene *FSP1* and the iPSC specific gene *NANOG*. Transiently reprogrammed samples expressed these genes at levels similar to control fibroblasts. Bars represent the mean and error bars the standard deviation. Samples transiently reprogrammed for 13, 15 or 17 days were pooled. The number of samples in each group is indicated in brackets. (F) Principal component analysis of transient reprogramming (magenta points) and reference reprogramming sample methylomes (light blue to dark blue, data from Banovich et al (2018), Ohnuki et al (2014) and our novel Sendai reprogramming dataset). Reference samples formed a reprogramming trajectory along PC1. Transient reprogramming samples moved along this trajectory with continued exposure to doxycycline (light magenta points) and returned to the beginning of the trajectory after withdrawal of doxycycline (magenta points). Control samples (yellow and orange points) remained at the beginning of the trajectory throughout the experiment. (G) Mean DNA methylation levels across the fibroblast specific gene *FSP1* and the iPSC specific gene *POU5F1* (encoding OCT4). Transiently reprogrammed samples had methylation profiles across these genes that resemble those found in fibroblasts. Grey bars and black bars indicate the locations of Ensembl annotated promoters and genes, respectively. Samples transiently reprogrammed for 10, 13, 15 or 17 days were pooled for visualisation purposes. The number of samples in each group is indicated in brackets.

After reprogramming for 10-17 days, we found the fibroblasts had undergone dramatic changes in morphology. Upon visual inspection using a light microscope it appeared that the cells had undergone a mesenchymal to epithelial like transition and were forming colony structures that progressively became larger with longer periods of reprogramming, consistent with the emergence of the early pluripotency marker SSEA4. After sorting the cells and culturing in the absence of doxycycline, we found they were able to return to their initial fibroblast morphology, showing that morphological reversion is possible even after 17 days of reprogramming (Figure 1C). We investigated further the identity of the cells after this MPTR by conducting DNA methylation array analysis and RNA sequencing to examine their methylomes and transcriptomes, respectively. We included published reprogramming datasets in our analysis as well as a novel reprogramming dataset that we generated based on Sendai virus delivery of the Yamanaka factors to act as a reference (Banovich et al., 2018; Fleischer et al., 2018; Ohnuki et al., 2014). Principal component analysis using expression values of all genes in the transcriptome separated cells based on extent of reprogramming and the reference datasets formed a reprogramming trajectory along PC1 (Figure 1D). Transiently reprogrammed samples clustered at the beginning of this trajectory showing that these samples transcriptionally resemble fibroblasts rather than reprogramming intermediates or iPSCs (Figure 1D). As examples, transiently reprogrammed cells did not express the pluripotency marker *NANOG* and expressed high levels of the fibroblast marker *FSP1* (Figure 1E). Similarly, principal component analysis of the methylomes separated cells based on extent of reprogramming and the reference datasets formed a reprogramming trajectory. Indeed, intermediate samples from our transient reprogramming experiment (collected after the reprogramming phase but before the reversion phase) clustered along this reprogramming trajectory (Figure 1F), showing that cells move epigenetically towards pluripotency consistent with the loss of the fibroblast surface marker CD13 and gain of the iPSC surface marker SSEA4. Notably, the transiently reprogrammed samples returned back to the start of this trajectory (with the reference fibroblast samples) revealing that they epigenetically resembled fibroblasts once again. We found typical regions that change during reprogramming returned to fibroblast-like states (Takahashi et al., 2007), such as the promoter of *OCT4* being hypermethylated and the promoter of *FSP1* being hypomethylated in our transiently reprogrammed cells (Figure 1G). Together, these data demonstrate that fibroblasts can be transiently reprogrammed to the maturation phase and then revert to a state that is morphologically, epigenetically and transcriptionally similar to the starting cell identity. To our knowledge, this is the first method for maturation phase transient reprogramming, where Yamanaka factors are transiently expressed up to the maturation phase of reprogramming before expression of the factors is abolished.

### Enhancer methylation memory persists at the intermediate stage of transient reprogramming

Though transiently reprogrammed fibroblasts temporarily lost their cell identity (becoming SSEA4 positive) they were able to reacquire it once the reprogramming factors were removed, suggesting that they retained memory of their initial cell identity. We hypothesised this may be in the form of epigenetic memory and so examined DNA methylation levels at promoters and enhancers during the reprogramming process. We used the Ensembl Regulatory Build to obtain the locations of promoter and enhancer elements as well as their activity status in dermal fibroblasts and iPSCs. We focussed on promoters and enhancers that are active in fibroblasts but are no longer active in iPSCs and classified these as fibroblast-specific promoters and enhancers (Figure 2A, B, bottom panels). Overall, we found that DNA methylation levels were relatively low at fibroblast-specific promoters, even in iPSCs (Figure 2A), except for a small subset of promoters (14 out of 479) that became hypermethylated during reprogramming but were still lowly methylated at the intermediate stage of transient reprogramming. These promoters may confer some memory of initial cell type, but they were not obviously related to fibroblast function (Figure 2A). Thus, promoter DNA methylation is unlikely to be the source of substantial memory of fibroblast identity. On the contrary, we found that DNA methylation was relatively dynamic at fibroblast-specific enhancers. Approximately half of all fibroblast-specific enhancers (2351 out of 4204) gain DNA methylation during iPSC reprogramming. However, even at day 17 of the reprogramming process (the longest transient reprogramming intermediate tested here), these enhancers still remained demethylated (Figure 2B). We also compared the methylation at these enhancers in our intermediate and final samples to reference datasets using PCA. Samples remain fibroblast-like at these enhancers until around day 20 of reprogramming when the stabilisation phase begins (Figure 2C), unlike the methylome as a whole (Figure 1F). Overall therefore, these enhancers are likely to remain poised during transient reprogramming and as a result may act as the source of memory for initial cell identity during a time when the somatic transcriptional program is otherwise repressed and somatic proteins such as CD13 are lost (David and Polo, 2014).

**Figure 2.**
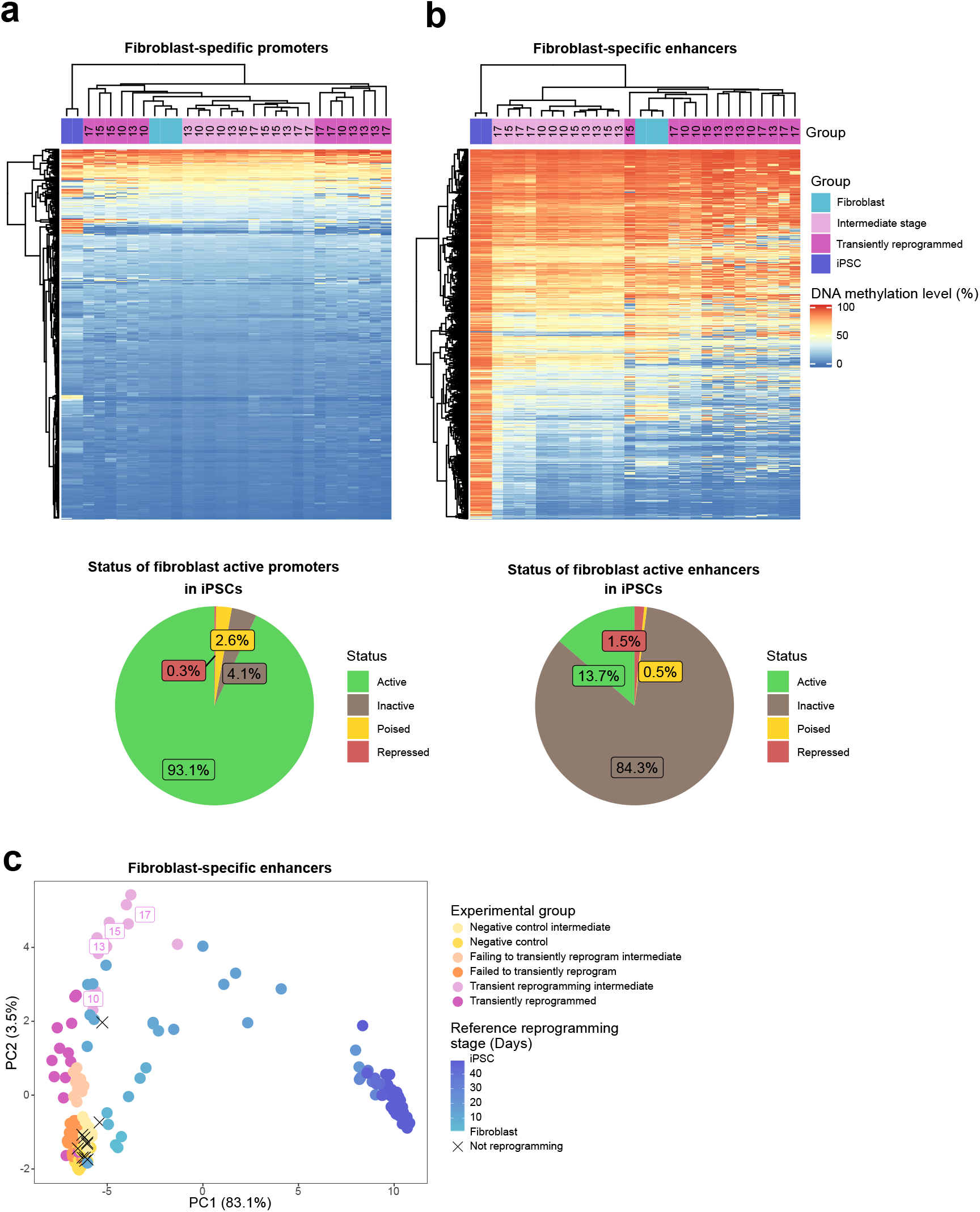
Epigenetic memory at enhancers may allow cells to return to their initial identity. (A) Heatmap (top panel) examining the DNA methylation levels of fibroblast-specific promoters in cells before (light blue group), during (light magenta group, transient reprogramming intermediate cells defined in Figure 1B) and after (magenta group, transiently reprogrammed fibroblasts defined in Figure 1B) transient reprogramming as well as in iPSCs (dark blue group). Each sample was plotted as a single column, whether reprogrammed for 10, 13, 15, or 17 days. Fibroblast promoters largely remain demethylated during complete reprogramming and at the intermediate stages of transient reprogramming. Fibroblast-specific promoters were defined as promoters that are active in fibroblasts but no longer active in iPSCs based on Ensembl regulatory build annotations (become inactive, poised or repressed). This occurs in only a small subset of promoters that are active in fibroblasts as shown in the pie chart (545 out of 7910; bottom panel). (B) Heatmap (top panel) examining the DNA methylation levels of fibroblast-specific enhancers in cells before (light blue group), during (light magenta group) and after (magenta group) transient reprogramming as well as in iPSCs (dark blue group). Each sample was plotted as a single column, whether reprogrammed for 10, 13, 15, or 17 days. Fibroblast enhancers became hypermethylated during complete reprogramming but were still demethylated at the intermediate stages of transient reprogramming. Fibroblast-specific enhancers were defined as enhancers that are active in fibroblasts but no longer active in iPSCs (become inactive, poised or repressed) based on Ensembl regulatory build annotations. This occurs to the majority of enhancers that are active in fibroblasts as shown in the pie chart (13831 out of 16026; bottom panel). (C) Principal component analysis of methylation data from transient reprogramming (magenta points) and reference reprogramming samples at fibroblast-specific enhancers (light blue to dark blue, data from Banovich et al (2018), Ohnuki et al (2014) and our novel Sendai reprogramming dataset). Reference samples formed a reprogramming trajectory along PC1. Transient reprogramming intermediate samples and transiently reprogrammed samples cluster closely with fibroblasts along PC1.

### Transient reprogramming reverses age-associated changes in the transcriptome

To investigate the effects of MPTR on the ageing transcriptome, we trained a transcription age-predictor using random forest regression on published fibroblast RNA-seq data from donors aged 1-94 years old (Fleischer et al., 2018). The transcription age predictor was trained on transformed age, similar to the Horvath epigenetic clock, to account for the accelerated ageing rate during childhood and adolescence (Horvath, 2013). The final transcription age predictor had a median absolute error of 13.5 years, this error being higher than that of the epigenetic clock consistent with previous transcription age predictors (Fleischer et al., 2018; Peters et al., 2015). Using our predictor, we found that transient reprogramming reduced transcription age by approximately 30 years and that all MPTR reprogramming times investigated were capable of reducing transcription age to a similar extent. We also observed a moderate reduction in transcription age in cells that failed to transiently reprogram (SSEA4 negative at the intermediate timepoint), suggesting expression of the reprogramming factors alone is capable of rejuvenating some aspects of the transcriptome (Figure 3A). This reduction in transcription age is substantially greater than that recently achieved by transient transfection of Yamanaka factors (Sarkar et al., 2020), which was by approximately 10 years according to our transcription age predictor (Supplementary Figure 2A), consistent with our approach of reprogramming further into the maturation phase rather than only up to the end of the initiation phase. To examine this transcriptomic rejuvenation in more detail, we identified genes that significantly correlated with age in the reference fibroblast ageing dataset (Fleischer et al., 2018) and used these genes to carry out principal component analysis (3707 genes). The samples primarily separated by age and reference fibroblast samples formed an ageing trajectory. The transiently reprogrammed samples clustered closer to the young fibroblasts along PC1 than the negative control samples, further supporting the findings of our transcription age predictor that MPTR rejuvenates the transcriptome (Figure 3B).

**Figure 3.**
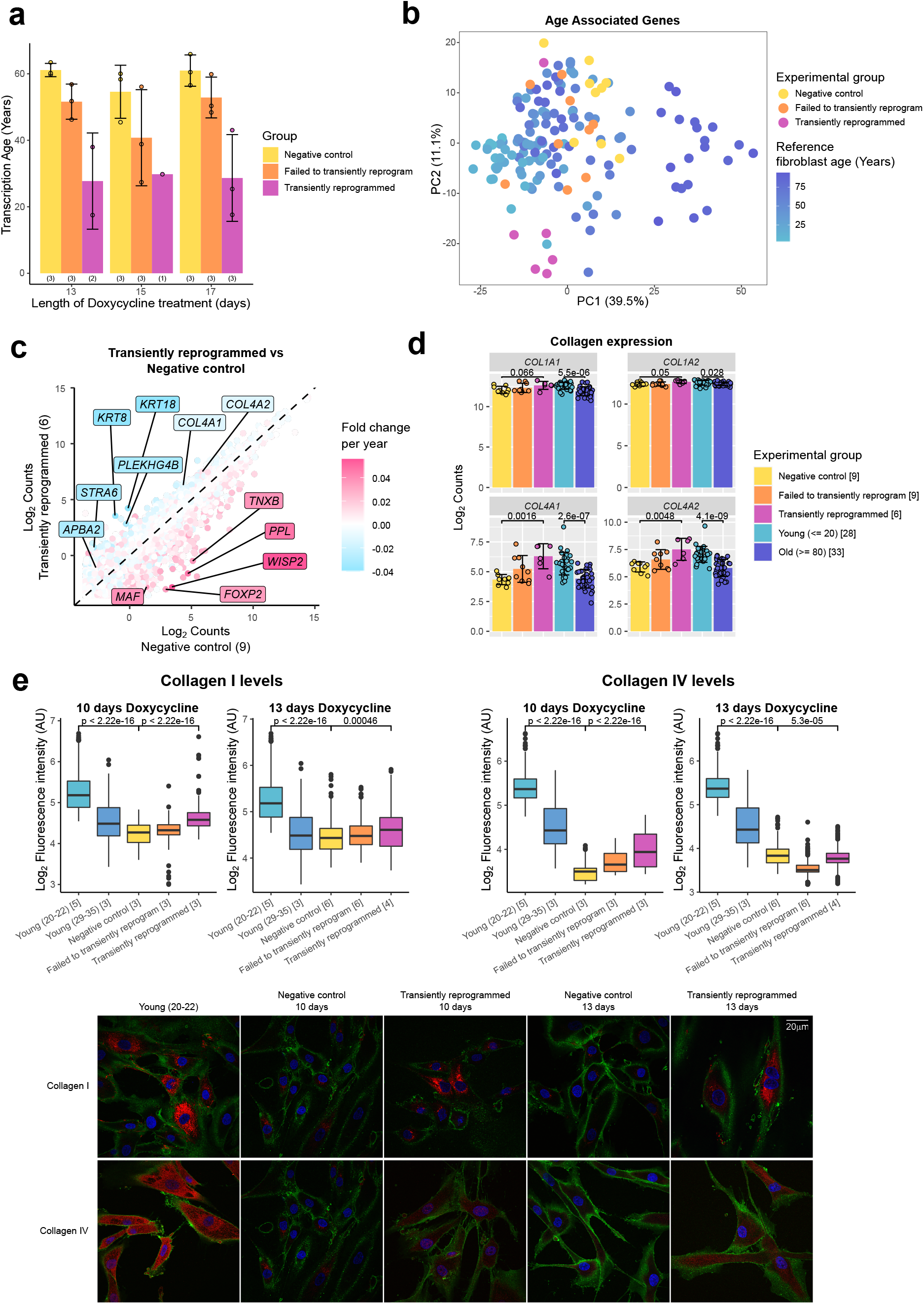
Transient reprogramming reverses age-associated changes in the transcriptome. (A) Mean transcription age calculated using a custom transcriptome clock (median absolute error=13.5 years) for negative control samples (yellow), samples that expressed OSKM but failed to reprogram based on cell surface markers (orange) and cells that were successfully transiently reprogrammed (magenta) as described in Figure 1B for 13, 15 or 17 days. The number of samples in each group is indicated in brackets. Bars represent the mean and error bars the standard deviation. (B) Principal component analysis of fibroblast ageing-associated gene expression levels in transient reprogramming (magenta) and reference ageing fibroblast samples (light blue-dark blue). Reference samples formed an ageing trajectory along PC1. Transiently reprogrammed samples located closer to young fibroblasts than negative control samples did, suggesting they were transcriptionally younger. (C) The mean expression levels of all genes in transiently reprogrammed samples compared to those in negative control samples. In addition, genes have been colour coded by their expression change with age. Genes that upregulate with age were downregulated with transient reprogramming and genes that downregulate with age were upregulated with transient reprogramming. Notable example genes have been highlighted. The number of samples in each group is indicated in brackets. (D) The expression levels of collagen genes that were restored to youthful levels after transient reprogramming. Samples transiently reprogrammed for 13, 15 and 17 days were pooled. Bars represent the mean and error bars the standard deviation. The number of samples in each group is indicated in square brackets. Significance was calculated with the Mann-Whitney U test. (E) Boxplots of the protein levels of Collagen I and IV in individual cells after transient reprogramming for 10 or 13 days calculated based on fluorescence intensity within segmented cells following immunofluorescence staining. Boxes represent upper and lower quartiles and central lines the median. The protein levels of Collagen I and IV increased after transient reprogramming. The number of samples in each group is indicated in square brackets. Representative images are included (bottom panel). CD44 is coloured in green, Collagen I and IV are coloured in red and DAPI staining is coloured in blue. Significance was calculated with the Mann-Whitney U test.

Next, we looked at the whole transcriptome and overlaid the ageing expression change onto the expression change following transient reprogramming. As expected, we observed an overall reversal of the ageing trends, with genes upregulated during ageing being downregulated following transient reprogramming and genes downregulated during ageing being upregulated following transient reprogramming (Figure 3C, Supplementary Figure 2B). Notably, structural proteins downregulated with age that were upregulated upon transient reprogramming included the cytokeratins 8 and 18 as well as subunits of collagen IV.

The production of collagens is a major function of fibroblasts (Humphrey et al., 2014), thus we examined the expression of all collagen genes during fibroblast ageing and during transient reprogramming (Figure 3D). As shown previously (Lago and Puzzi, 2019; Varani et al., 2006), we found collagen I and IV were downregulated with age, with collagen IV demonstrating a more dramatic reduction. Notably the expression of both genes was restored to youthful levels after transient reprogramming (Figure 3D). We then assessed by immunofluorescence whether this increased mRNA expression resulted in increased protein levels and indeed found that transient reprogramming resulted in an increase in Collagen I and IV protein towards more youthful levels (Figure 3E). Our data show that transient reprogramming followed by reversion can rejuvenate fibroblasts both transcriptionally and at the protein level, at least based on collagen production. This indicates that our rejuvenation protocol can, in principle, restore youthful functionality in human cells.

### Optimal transient reprogramming can reverse age-associated changes in the epigenome

After finding evidence of transcriptomic rejuvenation, we sought to determine whether there were also aspects of rejuvenation in the epigenome. We initially examined global levels of H3K9me3 by immunofluorescence. H3K9me3 is a histone modification associated with heterochromatin that has been previously shown to be reduced globally with age in a number of organisms (Ni et al., 2012), including in human fibroblasts (O’Sullivan et al., 2010; Scaffidi and Misteli, 2006). We were able to confirm this observation and found that MPTR was able to substantially reverse this age-associated reduction back to a level comparable with fibroblasts from younger donors (with a mean age of 33 years old). Both 10 and 13 days of transient reprogramming increased global levels of H3K9me3 suggesting that this epigenetic mark, similar to the transcriptome, has a relatively broad window for rejuvenation by transient reprogramming. We also observed a slight increase in H3K9me3 levels in cells that failed to transiently reprogram, suggesting that expression of the reprogramming factors alone is capable of partially restoring this epigenetic mark (Figure 4A), as was observed for our transcriptome-based age-predictor (Figure 3A). The magnitude of rejuvenation in H3K9me3 levels in our transiently reprogrammed cells is similar to that observed from initiation phase transient reprogramming (Sarkar et al., 2020).

**Figure 4.**
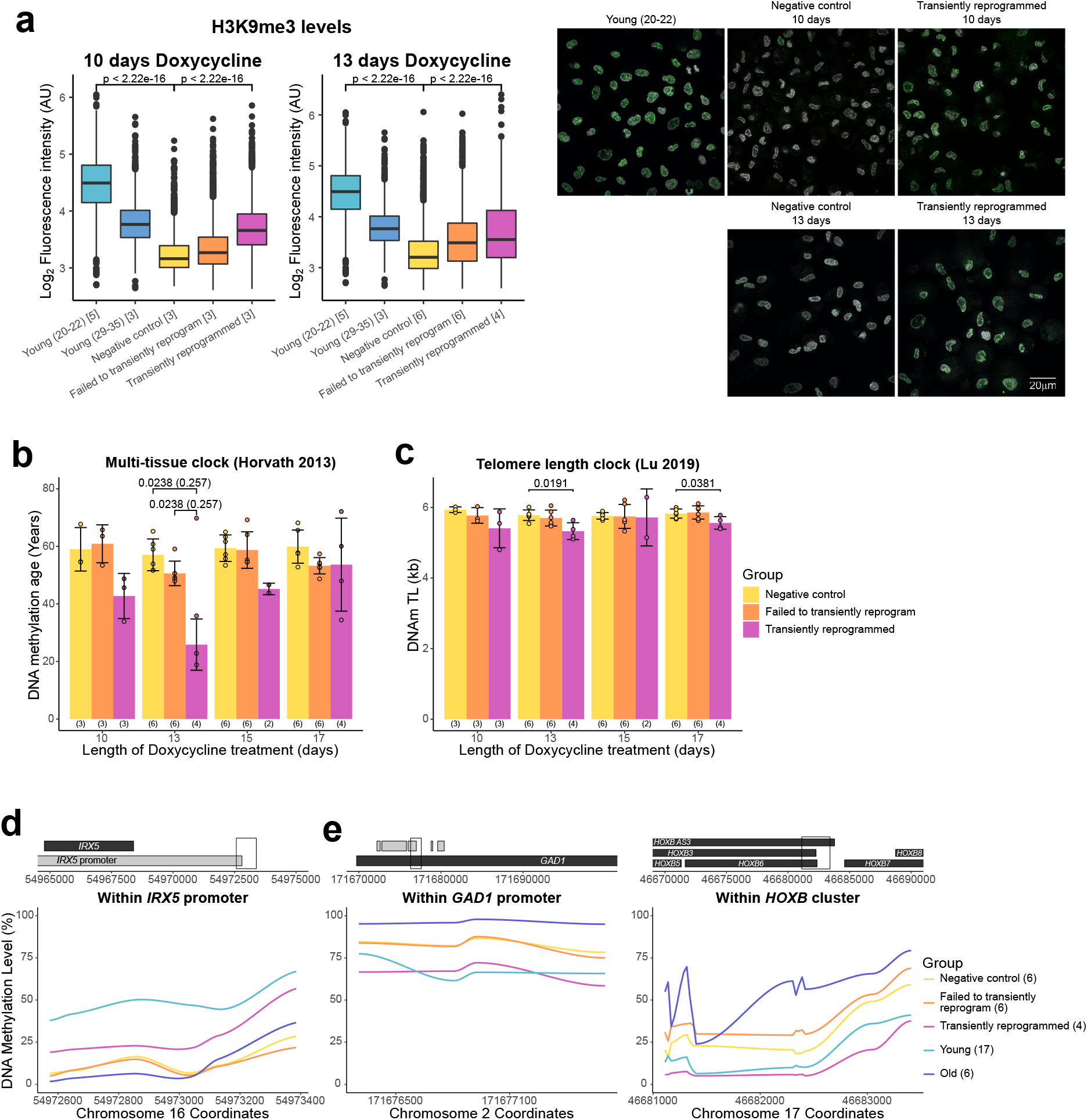
Optimal transient reprogramming can reverse age-associated changes in the epigenome. (A) Boxplots of the levels of H3K9me3 in individual cells calculated based on fluorescence intensity within nuclei (segmented using DAPI). The levels of H3K9me3 were found to decrease with age and increase after transient reprogramming for 10 or 13 days. Boxes represent upper and lower quartiles and central lines the median. The number of samples in each group is indicated in square brackets. Representative images are included (right panel). H3K9me3 is coloured in green and DAPI staining is coloured in greyscale. Significance was calculated with the Mann-Whitney U test. (B) Mean DNA methylation age of samples after transient reprogramming calculated using the multi-tissue clock (Horvath, 2013). DNA methylation age substantially reduced after 13 days of transient reprogramming. Shorter and longer lengths of transient reprogramming led to smaller reductions in DNA methylation age. Bars represent the mean and error bars represent the standard deviation. The outlier in the 13 days of transient reprogramming group was excluded from calculation of the mean and standard deviation. Significance was calculated with the Mann-Whitney U test with (in brackets) and without the outlier. The number of samples in each group is indicated in brackets beneath the bars. (C) Mean telomere length of samples after transient reprogramming calculated using the telomere length clock (Lu et al., 2019). Telomere length either did not change or was slightly reduced after transient reprogramming. Bars represent the mean and error bars represent the standard deviation. Significance was calculated with the Mann-Whitney U test. (D) Mean DNA methylation levels across a rejuvenated age-hypomethylated region. This region is found within the *IRX5* promoter. Samples transiently reprogrammed for 10, 13, 15 and 17 days were pooled for visualisation purposes. The number of samples in each group is indicated in brackets. (E) Mean DNA methylation levels across rejuvenated age-hypermethylated regions. These regions are found within the *GAD1* promoter and *HOXB* locus. Samples transiently reprogrammed for 10, 13, 15 and 17 days were pooled for visualisation purposes. The number of samples in each group is indicated in brackets.

We next applied the epigenetic clock, a multi-tissue age predictor that predicts age based on the DNA methylation levels at 353 CpG sites (Horvath, 2013), to our data. Notably, with 13 days of transient reprogramming we observed a substantial reduction of the median DNA methylation age – by approximately 30 years, quantitatively the same rejuvenation as we saw in the transcriptome (Figure 4B). A shorter period of transient reprogramming (10 days) resulted in a smaller reduction of DNA methylation age, consistent with our results profiling DNA methylation age throughout the reprogramming process, where DNA methylation age gradually reduced throughout the maturation phase (Figure 1A). Surprisingly, we also observed a smaller reduction in DNA methylation age with longer transient reprogramming times, suggesting that some aspects of the observed epigenetic rejuvenation are lost during the reversion phase of our MPTR protocol. Potentially, extended reprogramming (for 15 or 17 days) may make reversion more difficult and result in cellular stresses that ‘re-age’ the methylome during the process. Similar results were obtained using a different methylation age predictor, the skin and blood clock (Horvath et al., 2018)(Supplementary Figure 3A).

Telomeres are protective structures at the ends of chromosomes that consist of repetitive sequences. Telomere length decreases with age due to cell replication in the absence of telomerase enzymes and is restored upon complete iPSC reprogramming (Lapasset et al., 2011). To investigate the effect of transient reprogramming on telomere length we used the telomere length clock, which predicts telomere length based on the methylation levels at 140 CpG sites (Lu et al., 2019). We found that MPTR does not affect telomere length and, in some cases, slightly reduces it (Figure 4C). This is consistent with our results profiling telomere length throughout complete reprogramming using our doxycycline inducible system, where telomere length did not increase until the stabilisation phase (Supplementary Figure 3B).

Next, we investigated the locations of the rejuvenated CpG sites and found that most were individual sites spread across the genome (Supplementary Figure 3C). Some of these individual CpG sites may be part of larger regions of rejuvenated methylation, which we are unable to fully detect due to the targeted nature of DNA methylation array profiling, however, there were a few small clusters of rejuvenated CpG sites detected. We found that a small region in the *IRX5* promoter becomes demethylated with age and transient reprogramming is able to partially remethylate this region (Figure 4D). IRX5 is involved in embryonic development so demethylation of its promoter with age may lead to inappropriate expression (Cheng et al., 2005; Costantini et al., 2005). We also found two regions that become hypermethylated with age and are demethylated by transient reprogramming (Figure 4E). One of these regions is in the *GAD1* promoter; encoding an enzyme that catalyses the conversion of gamma-aminobutyric acid into glutamic acid (Bu et al., 1992). The other region is within the *HOXB* locus, involved in anterior-posterior patterning during development (Pearson et al., 2005). Overall, our data demonstrate that transient reprogramming for 13 days (but apparently not for longer or shorter periods) represents a ‘sweet spot’ that facilitates partial rejuvenation of the methylome, reducing epigenetic age by approximately 30 years.

## Discussion

Here we have developed a novel method, maturation phase transient reprogramming (MPTR), where Yamanaka factors are ectopically expressed until the maturation phase of reprogramming is reached, and their induction is then withdrawn. MPTR rejuvenates all measured molecular hallmarks of ageing robustly and substantially, including the transcriptome, epigenome, and functional protein expression. Previous attempts at transient reprogramming have been restricted to the initiation phase in order to conserve initial cell identity (Lu et al., 2020; Ocampo et al., 2016; Sarkar et al., 2020). This is a valid concern as fully reprogrammed iPSCs can be difficult to differentiate into mature adult cells and instead these differentiated cells often resemble their foetal counterparts (Hrvatin et al., 2014). With our approach, cells temporarily lose their cell identity as they enter the maturation phase but, importantly, reacquire their initial somatic fate when the reprogramming factors are withdrawn. This may be the result of persisting epigenetic memory at enhancers (Jadhav et al., 2019), which notably we find is not erased until the stabilisation phase. With our method employing longer periods of reprogramming, we observed robust and substantial rejuvenation of the whole transcriptome as well as aspects of the epigenome, with many features becoming approximately 30 years younger. This extent of rejuvenation is substantially greater than what has been observed previously for transient reprogramming approaches that reprogram within the initiation phase. The methylome appears to require longer reprogramming to substantially rejuvenate and consequently, previous work using shorter lengths of reprogramming has resulted in modest amounts of rejuvenation of the methylome (Lu et al., 2020; Sarkar et al., 2020).

Interestingly, we also observed transcriptomic rejuvenation in genes with non-fibroblast functions. In particular, the age-associated downregulation of *APBA2* and the age-associated upregulation of *MAF* were reversed (Figure 3C). APBA2 stabilises amyloid precursor protein, which plays a key role in the development of Alzheimer’s disease (Araki et al., 2003). MAF regulates the development of embryonic lens fibre cells, and defects in this gene lead to the development of cataracts, which are a frequent complication in older age (Ring et al., 2000). These observations may signal the potential of MPTR to promote more general rejuvenation signatures that could be relevant for other cell types such as neurons. It will be interesting to determine if MPTR-induced rejuvenation is possible in other cell types, which could help us understand and potentially treat age-related diseases such as Alzheimer’s disease and cataracts.

Overall, our results demonstrate that substantial rejuvenation is possible without acquiring stable pluripotency and suggest the exciting concept that the rejuvenation program may be separable from the pluripotency program. Future studies are warranted to determine the extent to which these two programs can be separated and could lead to discovery of novel targets that promote rejuvenation without the need for iPSC reprogramming.

## Methods

### Plasmids and lentivirus production

The doxycycline inducible polycistronic reprogramming vector was generated by cloning a GFP-IRES sequence downstream of the tetracycline response element in the backbone FUW-tetO-hOKMS (Addgene 51543). This vector was used in combination with FUW-M2rtTA (Addgene 20342). Viral particles were generated by transfecting HEK293T cells with the packaging plasmids pMD2.G (Addgene 12259) and psPAX2 (Addgene 12260) and either FUW-tetO-GFP-hOKMS or FUW-M2rtTA.

### iPSC reprogramming

Dermal fibroblasts from middle age donors (38-53 years old) were purchased from Lonza. For lentiviral iPSC reprogramming, fibroblasts were expanded in fibroblast medium (DMEM-F12, 10% FBS, 1X Glutamax, 1X MEM-NEAA, 1X beta-mercaptoethanol, 0.2X Penicillin/Streptomycin, 16 ng/ml FGF2) before being spinfected with tetO-GFP-hOKMS and M2rtTA lentiviruses, where 10% virus supernatant and 8 µg/ml polybrene was added to the cells before centrifugation at 1000 rpm for 60 minutes at 32°C. Reprogramming was initiated 24 hours after lentiviral transduction by introducing doxycycline (2 µg/ml) to the media. Media was changed daily throughout the experiment subsequently. On day 2 of reprogramming, cells were flow sorted for viable GFP positive cells and then cultured on gelatine coated plates. On day 7 of reprogramming, cells were replated onto irradiated mouse embryonic fibroblasts (iMEFs) and on day 8 of reprogramming, the medium was switched to hES medium (DMEM-F12, 20% KSR, 1X Glutamax, 1X MEM-NEAA, 1X beta-mercaptoethanol, 0.2X Penicillin/Streptomycin, 8 ng/ml FGF2). For transient reprogramming, cells were flow sorted at day 10, 13, 15 or 17 of reprogramming for the CD13+ SSEA4- and CD13-SSEA4+ populations. These cells were then replated on iMEFs in fibroblast medium without doxycycline and subsequently maintained like fibroblasts without iMEFs. For complete reprogramming, colonies were picked on day 30 and transferred onto Vitronectin coated plates in E8 medium without doxycycline. Colonies were maintained as previously described (Milagre et al., 2017) and harvested at day 51 of reprogramming.

For Sendai virus iPSC reprogramming using CytoTune™-iPS 2.0 Sendai Reprogramming kit (Invitrogen), fibroblasts were reprogrammed as previously described (Milagre et al., 2017).

### Fluorescence-activated cell sorting (FACS) of reprogramming intermediates

Cells were pre-treated with 10 µM Y-27632 (STEMCELL technologies) for 1 hour. Cells were harvested using StemPro™ Accutase™ cell dissociation reagent and incubated with antibodies against CD13 (PE, 301704, Biolegend), SSEA4 (AF647, 330408, Biolegend) and CD90.2 (APC-Cy7, 105328, Biolegend) for 30 minutes. Cells were washed twice with 2% FBS in PBS and passed through a 50 µm filter to achieve a single cell suspension. Cells were stained with 1 µg/mL DAPI just prior to sorting. Single colour controls were used to perform compensation and gates were set based on the “negative control intermediate” samples. Cells were sorted with a BD FACSAria™ Fusion flow cytometer (BD Biosciences) and collected for either further culture or DNA/RNA extraction.

### DNA methylation array

Genomic DNA was extracted from cell samples with the DNeasy blood and tissue kit (Qiagen) by following the manufacturer’s instructions and including the optional RNase digestion step. For intermediate reprogramming stage samples, genomic DNA was extracted alongside the RNA with the AllPrep DNA/RNA mini kit (Qiagen). Genomic DNA samples were processed further at the Barts and the London Genome Centre and run on Infinium MethylationEPIC arrays (Illumina).

### RNA-Seq

RNA was extracted from cell samples with the RNeasy mini kit (Qiagen) by following the manufacturer’s instructions. For intermediate reprogramming stage samples and Sendai virus reprogrammed samples, RNA was extracted alongside the genomic DNA with the AllPrep DNA/RNA mini kit (Qiagen). RNA samples were DNase treated (Thermo Scientific) to remove contaminating DNA. RNA-Seq libraries were prepared at the Wellcome Sanger Institute and run on a HiSeq 2500 system (Illumina) for 50 bp single-end sequencing. For Sendai virus reprogrammed samples, libraries were prepared as previously described (Milagre et al., 2017), and run on a HiSeq 2500 (Illumina) for 75 bp paired-end sequencing.

### DNA methylation analysis

The array data was processed with the minfi R package and NOOB normalisation to generate beta values. DNA methylation age was calculated using the multi-tissue clock (Horvath, 2013) and the skin and blood clock (Horvath et al., 2018). Telomere length was calculated using the telomere length clock (Lu et al., 2019). Reference datasets for reprogramming fibroblasts and iPSCs were obtained from Ohnuki et al (2014) (GEO: GSE54848), Banovich et al (2018) (GEO: GSE110544) and Horvath et al (2018). In addition, the reference datasets included novel data examining the intermediate stages of dermal fibroblasts being reprogrammed with the CytoTune™-iPS 2.0 Sendai Reprogramming kit (Invitrogen).

### RNA-Seq analysis

Reads were trimmed with Trim Galore (version 0.6.2) and aligned to the human genome (GRCh38) with Hisat2 (version 2.1.0). Raw counts and log2 transformed counts were generated with Seqmonk (version 1.45.4). Reference datasets for fibroblasts and iPSCs were obtained from Fleischer et al (2018) (GEO: GSE113957) and Banovich et al (2018) (GEO: GSE107654). In addition, the reference datasets included novel data examining the intermediate stages of dermal fibroblasts being reprogrammed with the CytoTune™-iPS 2.0 Sendai Reprogramming kit (Invitrogen). Samples were carried forward for further analysis if they had a total read count of at least 500000 with at least 70% of the reads mapping to genes and at least 65% of the reads mapping to exons.

### Immunofluorescence and Imaging

Young control dermal fibroblasts were purchased from Lonza and the Coriell Institute (GM04505, GM04506, GM07525, GM07545 and AG09309). Antibody staining was performed as previously described (Santos et al., 2003) on cells grown on coverslips or cytospun onto coverslip after fixation with 2% PFA for 30 minutes at room temperature. Briefly, cells were permeabilised with 0.5% TritonX-100 in PBS for 1 hour; blocked with 1% BSA in 0.05% Tween20 in PBS (BS) for 1 hour; incubated overnight at 4°C with the appropriate primary antibody diluted in BS; followed by wash in BS and secondary antibody. All secondary antibodies were Alexa Fluor conjugated (Molecular Probes) diluted 1:1000 in BS and incubated for 30 minutes. Incubations were performed at room temperature, except where stated otherwise. DNA was counterstained with 5 μg/mL DAPI in PBS. Optical sections were captured with a Zeiss LSM780 microscope (63x oil-immersion objective). Fluorescence semi-quantification analysis was performed with Volocity 6.3 (Improvision). 3D rendering of z-stacks was used for semi-quantification of Collagen I and IV. Single middle optical sections were used for semi-quantification of H3K9me3. Antibodies and dilutions used are listed below:

Anti-H3K9me3; 07-442, Merck/ Millipore (1:500)

Anti-Collagen I; ab254113, Abcam (1:400)

Anti-Collagen IV; PA5-104508, Invitrogen (1:200)

Anti-CD44-BB515; 564587, BD Biosciences (1:400)

### Data analyses

Downstream analyses of RNA-seq and DNA methylation data were performed using R (version 4.0.2). Ggplot2 (version 3.3.2) was used to generate the bar charts, boxplots, line plots, pie charts, scatter plots and violin plots. Ggalluvial (version 0.12.2) was used to generate the alluvial plots. ComplexHeatmap (version 2.4.3) was used to generate the heatmaps. The combat function from the package sva (version 3.36.0) was used in figure 1F to batch correct the novel Sendai reprogramming dataset to the other datasets. The combat function was also used in figure 3 to batch correct the fibroblast ageing reference dataset (Fleischer et al., 2018) to our dataset. The transcription clock was trained on the batch corrected ageing reference dataset using the caret R package (Kuhn, 2008) and random forest regression with 10-fold cross validation. Rejuvenated CpG sites were found by comparing the methylation difference due to the age (calculated with the Horvath et al, 2018 dataset) to the methylation difference due to 13 days of transient reprogramming. CpG sites were classified as rejuvenated if they demonstrated a methylation difference of 10% over 40 years of ageing that was reversed by transient reprogramming.

## Author contributions

D.G. conceived the project, designed and performed the transient reprogramming experiments, carried out the data analysis and wrote the manuscript. A.P. helped design the transient reprogramming experiments and write the manuscript. F.S. performed the immunofluorescence experiments. I.H. provided bioinformatic support for the methylation analysis. T.M.S. provided helpful discussions for project design and helped with the design of the transient reprogramming experiments. I.M. provided helpful discussions for project design, designed and performed the Sendai reprogramming experiments and helped write the manuscript. W.R. provided helpful discussions for the project and for the interpretation of the data, and helped write the manuscript.

## Acknowledgements

We would like to thank all members of the Reik lab for helpful discussions. We would like to thank the bioinformatics facility at the Babraham Institute for processing the sequencing data, and the flow cytometry facility at the Babraham Institute for cell sorting. We would also like to thank the sequencing facilities at the Sanger Institute and the Bart’s and the London Genome Centre for sequencing and methylation array services respectively. This work was funded by the BBSRC. A.P. is supported by a Sir Henry Wellcome Fellowship (215912/Z/19/Z). W.R. is a consultant and shareholder of Cambridge Epigenetix. T.S. is CEO and shareholder of Chronomics.

## Figures

**Supplementary Figure 1.**
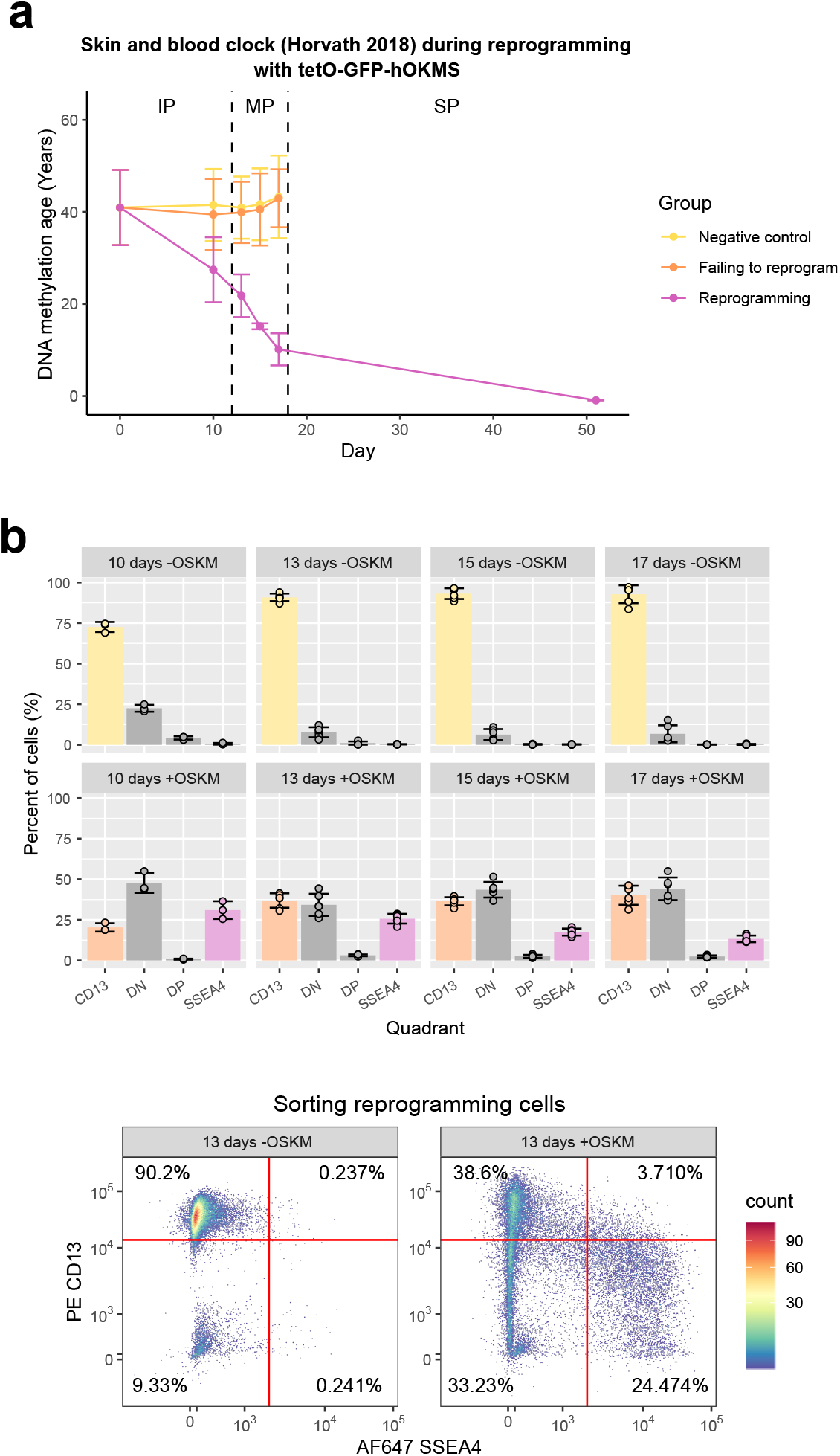
(A) Mean DNA methylation age (calculated using the skin and blood clock (Horvath et al., 2018)) throughout the reprogramming process where cells were transduced with our tetO-GFP-hOKMS vector and treated continuously with 2 µg/mL of doxycycline. Reprogramming is divided into three distinct phases: initiation phase (IP); maturation phase (MP) and stabilisation phase (SP). DNA methylation age decreased substantially during the maturation phase of reprogramming in cells that were successfully reprogramming (magenta line) but not in control cells (yellow and orange lines represent non-transduced cells and cells expressing hOKMS but failing to reprogram as indicated by cell surface markers, respectively). Points represent the mean and error bars the standard deviation. N = 3 biological replicates per condition, where fibroblasts were derived from different donors. N = 2 biological replicates for the iPSC timepoint (day 51). (B) The percentage of cells measured in each quadrant during the flow sort for successfully reprogramming cells (top panel). Cells were classified as CD13 only (CD13+ SSEA4-), double negative (“DN”, CD13-SSEA4-), double positive (“DP”, CD13+ SSEA4+) or SSEA4 only (CD13-SSEA4+). Cells that were collected are colour coded (light yellow = negative control intermediate, light orange = failed to transiently reprogram intermediate, light magenta = transient reprogramming intermediate). Bars represent the mean and error bars the standard deviation. Representative flow cytometry plots (bottom panel) show where gates were placed for determining presence/absence of surface markers.

**Supplementary Figure 2.**
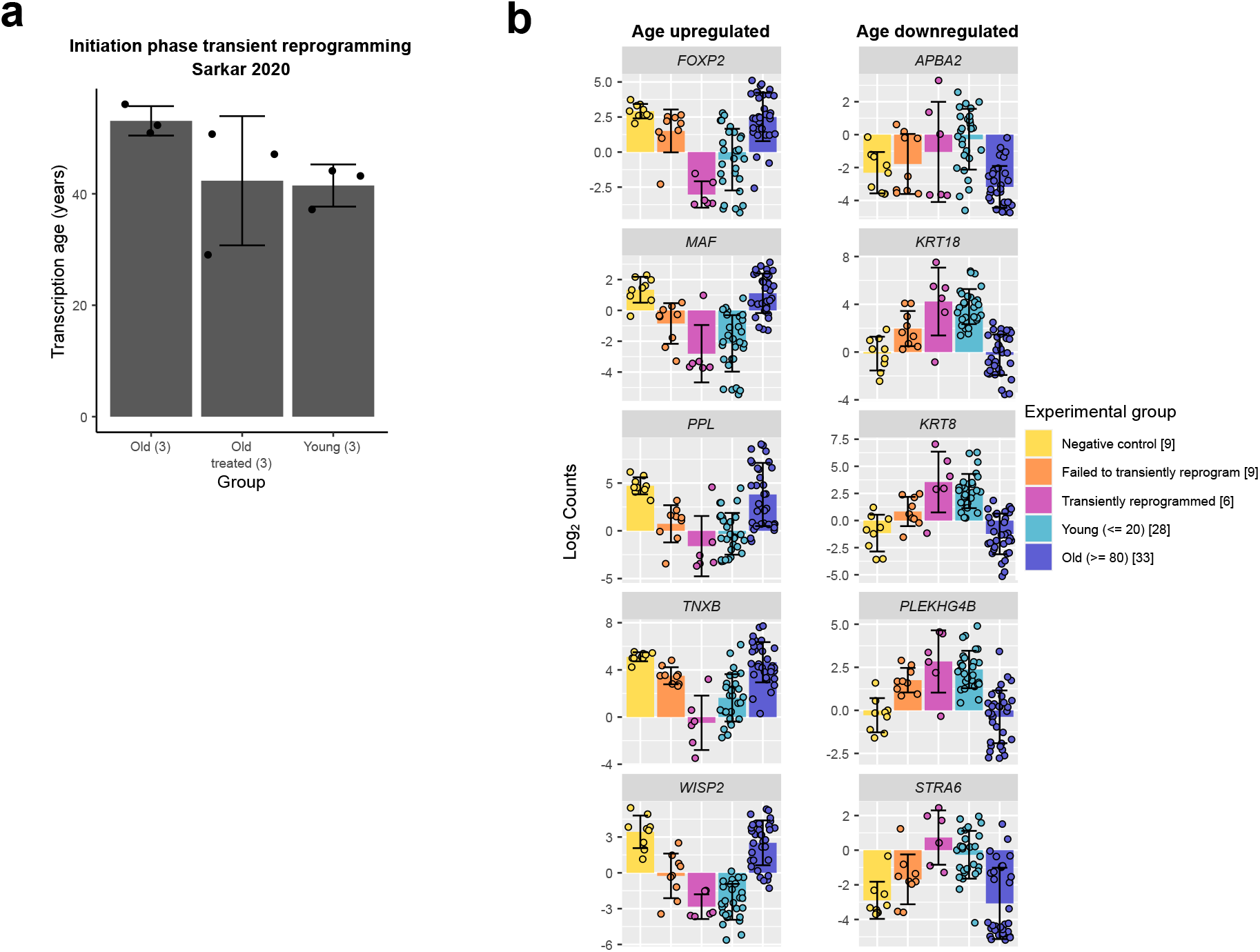
(A) Mean transcription age calculated using a custom transcriptome clock (error=13.5 years) for initiation phase transiently reprogrammed fibroblasts (Sarkar et al., 2020). The number of samples in each group is indicated in brackets. Bars represent the mean and error bars the standard deviation. (B) The expression levels of notable genes that were restored to youthful levels after transient reprogramming. Samples transiently reprogrammed for 13, 15 and 17 days were pooled. Bars represent the mean and error bars the standard deviation. The number of samples in each group is indicated in square brackets.

**Supplementary Figure 3.**
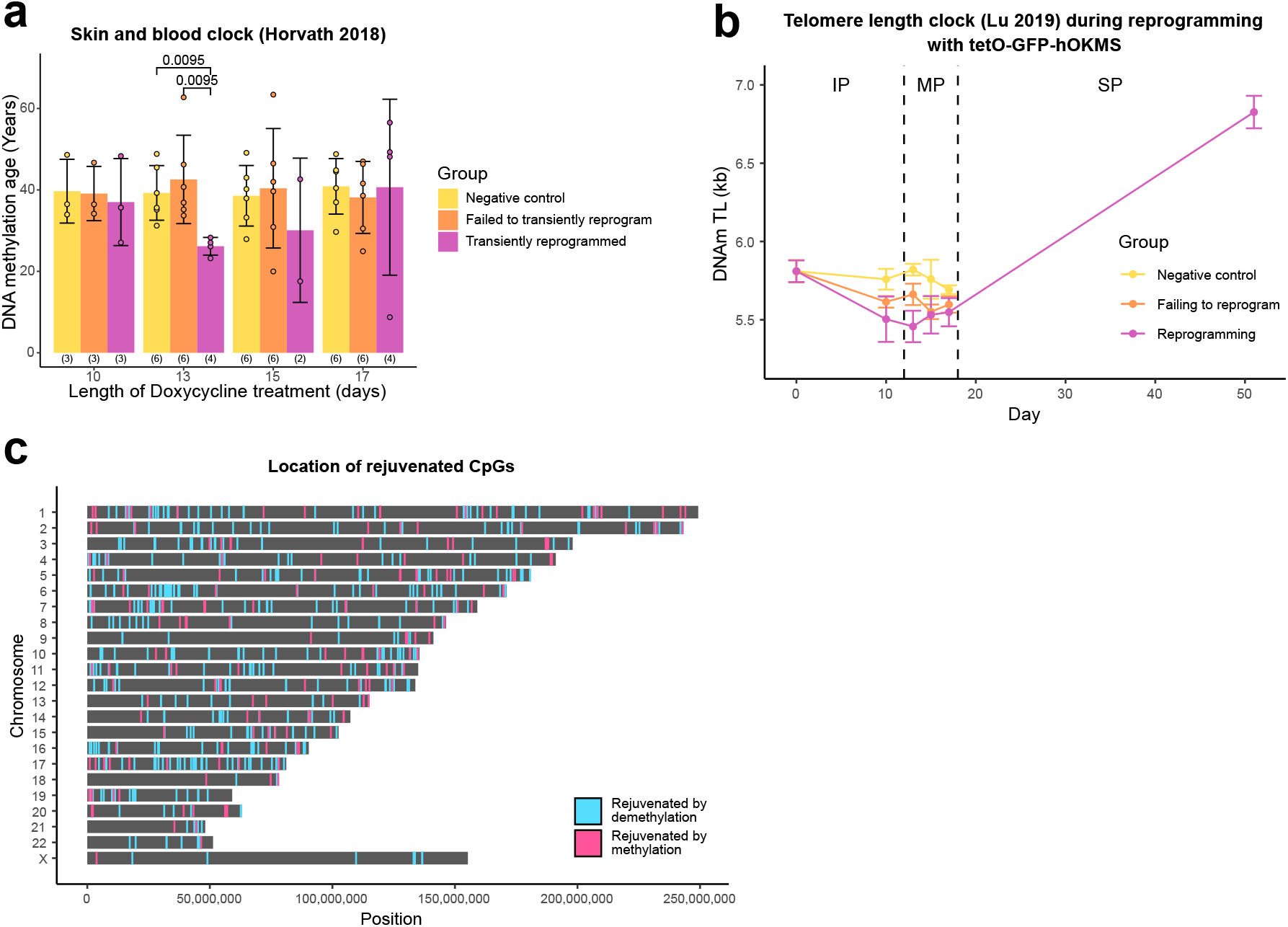
(A) Mean DNA methylation age of samples after transient reprogramming calculated using skin and blood clock (Horvath et al., 2018). DNA methylation age substantially reduced after 13 days of transient reprogramming. Shorter and longer lengths of transient reprogramming led to smaller reductions in DNA methylation age. Bars represent the mean and error bars represent the standard deviation. Significance was calculated with the Mann-Whitney U test. The number of samples in each group is indicated in brackets. (B) Mean telomere length (calculated using the telomere length clock (Lu et al., 2019)) throughout the reprogramming process where cells were transduced with our tetO-GFP-hOKMS vector and treated continuously with 2 µg/mL of doxycycline. Reprogramming is divided into three distinct phases: initiation phase (IP); maturation phase (MP) and stabilisation phase (SP). Telomere length decreases during the initiation and maturation phases and begins to increase during the stabilisation phase. Points represent the mean and error bars the standard deviation. N = 3 biological replicates per condition, where fibroblasts were derived from different donors. N = 2 biological replicates for the iPSC timepoint (day 51). (C) The location of rejuvenated CpG sites after 13 days of transient reprogramming. Sites rejuvenated by demethylation are coloured in blue and sites rejuvenated by methylation are coloured in pink.

